# Differential Analysis of Alternative Splicing Events in gene regions using Residual Neural Networks

**DOI:** 10.1101/2024.10.30.621059

**Authors:** Simone Ciccolella, Luca Denti, Jorge Avila Cartes, Gianluca Della Vedova, Yuri Pirola, Raffaella Rizzi, Paola Bonizzoni

**Affiliations:** DISCo, Università degli Studi di Milano-Bicocca, Milan, Italy

## Abstract

Several computational methods for the differential analysis of alternative splicing (AS) events among RNA-seq samples typically rely on estimating isoform-level gene expression. However, these approaches are often error-prone due to the interplay of individual AS events, which results in different isoforms with locally similar sequences. Moreover, methods based on isoform-level quantification usually need annotated transcripts.

In this work, we leverage the ability of deep learning networks to learn features from images, to propose `deepSpecas`, a novel method for event-based AS differential analysis between two RNA-seq samples. Our method does not rely on isoform abundance estimation, neither on a specific annotation. `deepSpecas` employs an image embedding scheme to represent the alignments of the two samples on the same region and utilizes a residual neural network to predict the AS events possibly expressed within that region. To our knowledge `deepSpecas` is the first deep learning approach for performing an event-based AS analysis of RNA-seq samples. To validate `deepSpecas`, we also address the lack of high quality AS benchmark datasets. For this purpose, we manually curated a set of regions exhibiting AS events. These regions were used for training our model and for comparing our method with state-of-the-art event-based AS analysis tools. Our results highlight that `deepSpecas` achieves higher precision at the expense of a small reduction in sensitivity.

The tool and the manually curated regions are available at https://github.com/sciccolella/deepSpecas.

## 1 Introduction

Alternative Splicing (AS) is the mechanism that allows the expression of multiple proteins from a single gene and it is involved both in physiological and pathological contexts [1, 2].

The emergence of Next-Generation Sequencing (NGS) technologies, with their high throughput, compared with pre-NGS technologies, brought forth the promise of revolutionizing gene expression analysis in two ways: (1) enabling the exploration of gene expression at the transcript level and (2) facilitating the discovery of novel genes and transcripts within the entire transcriptome [1].

However, early NGS technologies were severely limited in read length, which negatively affected their ability to differentiate repeats. This limitation is relevant in RNA-seq analyses since, due to AS, different isoforms usually share large portions of their sequence. As a consequence, *transcript quantification* is very complex, because RNA-seq reads cannot be unambiguously assigned to their originating transcript. Transcript quantification is typically followed by differential expression analysis, that aims at identifying the genes and/or transcripts whose expressions were significantly different between conditions. Differential expression analysis is important, since it enables to focus the attention only to those few genes or transcripts that, among tens of thousands, could potentially be involved in the expression of the phenotype under study. Many tools in the literature aim at quantifying transcripts. For example, StringTie [3], Cufflinks [4], Scripture [5] and IsoLasso [6] are transcript assembly tool also performing the quantification of the reconstructed sequences. Kallisto [7] and Salmon [8] perform transcript quantification taking as input a transcriptome and a set of RNA-Seq reads On the other hand, a less ambitious, but quite relevant task, is the detection of AS events that differentiate the transcripts expressed from two or more samples of RNA-seq. In this case, the focus is not on predicting transcripts whose expressions are significantly different between conditions, but instead predicting the AS events that differentiate two conditions. For this task, some alignment-based approaches have been proposed. For example, rMATS [9], SpliceSeq [10], Spladder [11], ASGAL [12], ESGq [13], and pantas [14] are designed to detect AS events (exon skipping, 5’ and 3’ competing sites, intron retention, etc.). Suppa2 [15], JunctionSeq [16] and MAJIQ [17] perform quantification at a splice junction level, while DexSeq [18] identifies differentially expressed exons. Obsrve that these methods start by aligning reads either to the genome or to the transcriptome and therefore they often need the support of an existing gene annotation.

We follow the research direction started by some state-of-art tools, exploiting deep learning (Convolutional Neural Networks CNN, in particular, Residual Neural Networks ResNet) to accomplish bioinformatics tasks such as variant calling, classifying or detecting splicing sites and gene expression analysis [19, 20]. For example, DeepVariant [21] identifies variants in NGS reads, SpliceAI [22] exploits a deep learning network to annotate input variants with their predicted effect on splicing, SpliceRover [23] and DeepSplicer [24] make splice site prediction, CI-SpliceAI [25] predicts disease causing splicing variants. We propose a novel method for detecting AS events in two RNA-Seq samples, leveraging on an image embedding schema representing alignments of the samples to the genome. As far as we know, this is the first method that uses a deep learning approach for detecting AS events expressed by two RNA-Seq samples, a description of the method is depicted in Figure 1. Our main goal is to avoid relying on a specific transcript annotation, that often constitutes a limitation, as it might be incomplete or even unavailable for contexts such as detecting alternative splicing in tumor cells [26], where we do not have a reliable annotation. We implemented our deep learning method in the tool `deepSpecas` which takes as input the alignments (to the genome) of two RNA-Seq samples and a list of genomic regions (*i*.*e*., genomic locations) in order to detect, for each specified region, the AS event (cassette exon, competing sites, intron retention, etc.) one sample expresses with respect to the other.

**Figure 1:**
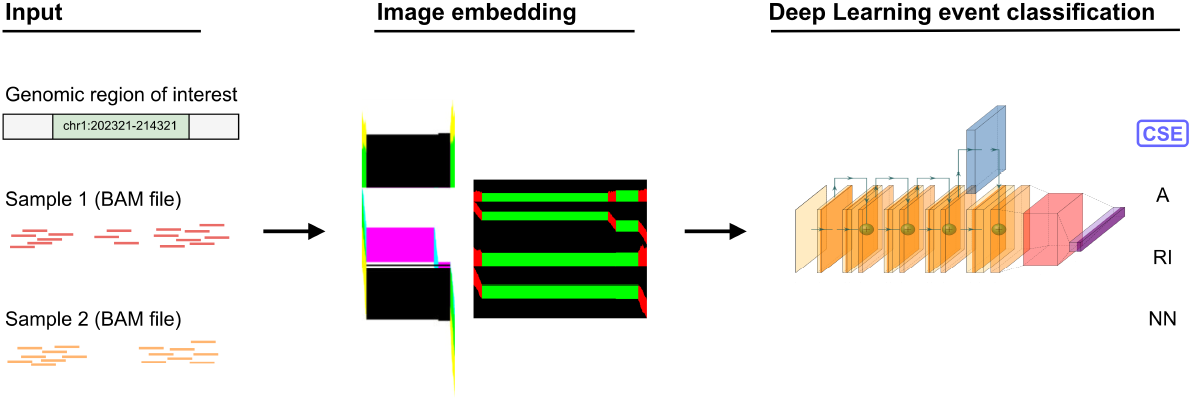
Graphical representation of `deepSpecas` framework. The tool requires as input a genomic region of interested, represented in standard forward notation, and a pair of samples (conditions) of splice-aligned reads in BAM format; `deepSpecas` proceeds to produce required image embedding of the two samples for the specified region and using the RNN model outputs the predicted differential AS events between the two samples, or *non-event* if none is expressed.

To validate the performance of `deepSpecas`, we also curated a comprehensive benchmark. Indeed, existing benchmark studies, which may help users select appropriate tools for their analyses, have several limitations. Firstly, these studies often use as ground truth only a small subset of well-studied genes or rely on simulated RNA-Seq data, where alternative splicing events are introduced randomly and encompass only few event types. We have evaluated `deepSpecas` with the classical ML accuracy metrics (precision, recall and F1-score), showing extremely high F1-scores (99%+) on (i) train-test-validation and (ii) with 5-fold cross-validation with 70:30 splits and (iii) very high accuracy on real dataset extracted by consensus of the two state-of-the-art tools `rMATS` and `SUPPA2`.

Our last contribution of this work is a detailed analysis of a real dataset, with a detailed and thorough examination of the AS events, consisting of both a visual inspection with IGV [27] and statistical examinations of the (several) predicted AS events that are imprecise or doubtful. Afterwards, we have manually curated a set of events that can be used by future works as an accurate dataset for benchmark of performance.

## 2 Methods

`deepSpecas` takes as input the alignments (given in BAM format) of two RNA-Seq samples (denoted as *Sample1* and *Sample2* in the rest of the paper) to the reference genome and a list of genomic regions, referred to as *regions of interest*, where potentially the two samples exhibit an AS event among the following: Cassette Exon/Exon Skipping (CSE), Alternative Acceptor/Donor sites (A) and Intron Retention (IR). `deepSpecas` exploits a Residual Neural Network (ResNet) to decide, for each input region of interest, whether the two samples exhibit a particular alternative splicing event (among the previous ones) relative to each other. For example, an exon included in *Sample1* but not in *Sample2*, or vice versa (Cassette Exon/Exon Skipping, CSE), a donor/acceptor competing site between an exon in *Sample1* and an exon in *Sample2* (A), or an exon of *Sample1* within which is retained an intron in *Sample2*, or vice versa (RI). The tool will report a *non-event* NN if no alternative event has been detected between the two samples.

The method used by `deepSpecas` is the following. Alignments (from the two samples) falling within a given region are encoded through an image (bi- or four-dimensional *data tensor* depending on the encoding), and an image for each region of interest is produced. The trained ResNet is fed with each image produced, thereby yielding the response for each one of the considered regions of interest.

Section 2.1 describes the types of images we use for encoding the alignments inside a given region. Section 2.2 describes the generation of the training images, while Secions 2.3 and 2.4 present the model architecture and the model training.

### 2.1 Encoding the regions of interest

In this work we explore the accuracy achieved by the ResNet using six different encodings (i.e., data tensors) of the read alignments on each region of interest. The encodings provide an image representation of the alignments in the region and mimic the commonly-used views of spliced alignments on genomic viewers such as IGV. In particular, the six encodings we propose are:

- **Cov:** the image encodes vertical bar plots representing the coverage (*y* - axis) across the genomic region (*x* -axis). Each pixel on the *x* -axis represents a single genomic position and the coverage at each position is the height (in pixels) of the bar. Two different coverages are computed and encoded as distinct channels of the image: the read coverage, *i*.*e*., the number of reads aligned on that position, and the *reference skip* coverage, *i*.*e*., the number of alignments that contain a N in their CIGAR string in that position (in other words, the number of alignments that “support” the presence of an intron in that position). The two coverages of each sample are represented by a single bar plot. The two bar plots of the two samples are vertically stacked. An example is shown in Figure 2a, in which read coverage uses the red channel and reference skip coverage the green channel.
- **Squish:** the image displays the read alignments of the two samples in the region, one sample on top of the other. Each read alignment is represented as a horizontal, 1 pixel high, line of the image. Alignments are ordered according to their starting position on the genome. Aligned positions and reference skips (*i*.*e*., Ns in the CIGAR string) are encoded on two separate channels. An example is shown in Figure 2b, in which aligned positions use the red channel and skipped positions the green channel.
- **Comb:** vertically stacks the two previous encodings, grouped by sample. In other words, the resulting image is the *Cov* encoding and *Squish* encoding of the first sample on top of the *Cov* encoding and *Squish* encoding of the second sample. An example is shown in Figure 2c.
- **Cov-4D:** contains the same information as *Cov* but the two samples are superimposed on separate channels instead of being vertically stacked, resulting in a 4-channel image (see Figure 3a).
- **Squish-4D:** contains the same information as *Squish* but the two samples are superimposed on separate channels instead of being vertically stacked, resulting in a 4-channel image (see Figure 3b).
- **Comb-4D:** contains the same information as *Comb* but the two samples are superimposed on separate channels instead of being vertically stacked, resulting in a 4-channel image (see Figure 3c).

**Figure 2:**
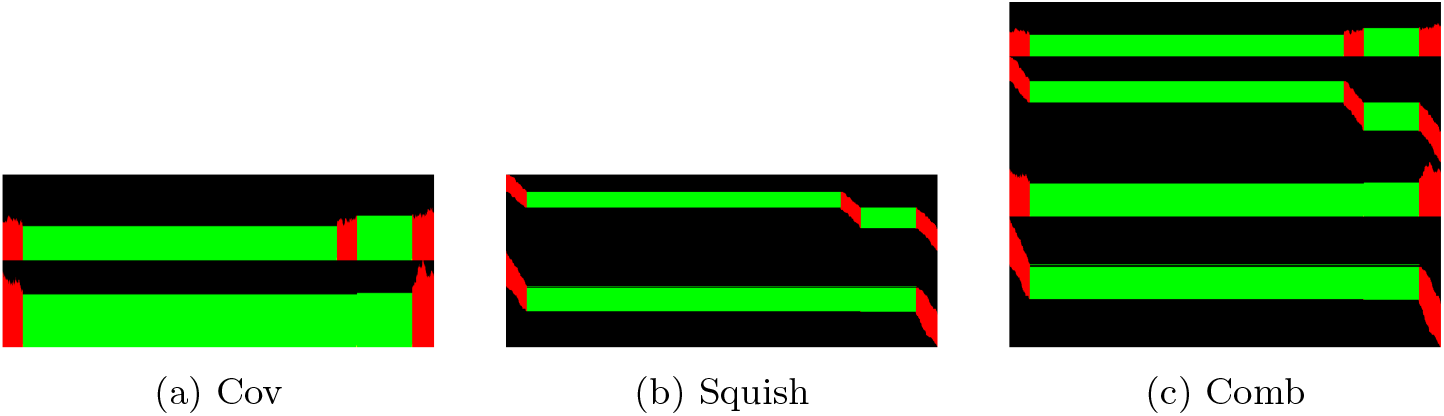
Three image representations of the read alignments of the same genomic region using 2-channel images. Red pixels represent *aligned* portions, green pixel *reference skips* and black pixel no information. This example region exhibits an *Exon Skipping/Cassette Exon* event.

**Figure 3:**
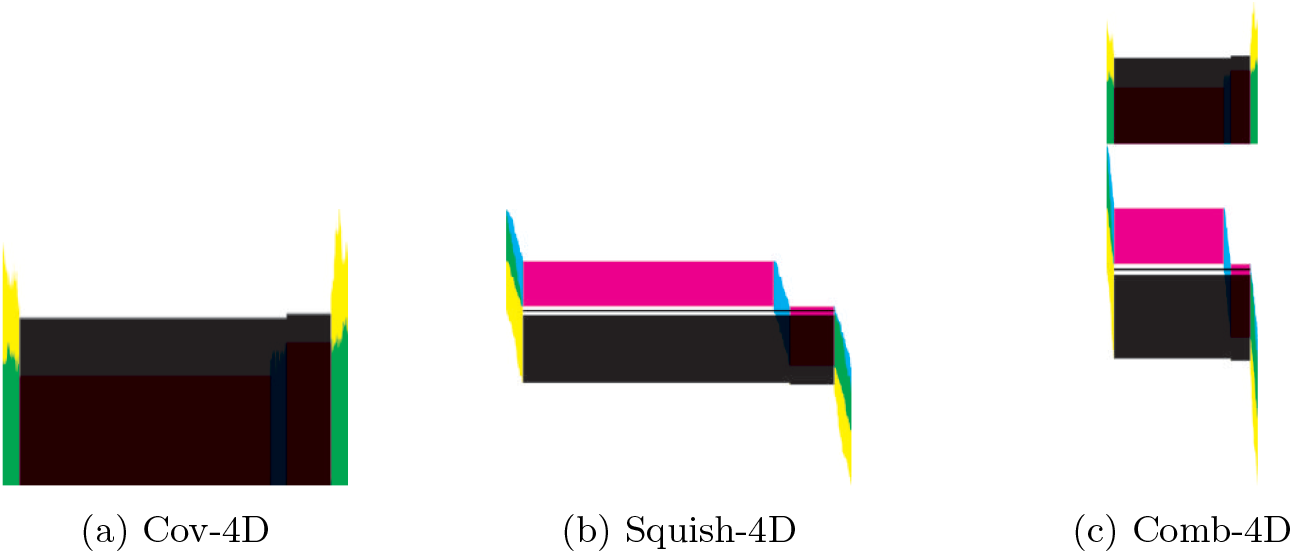
Three image representations of the read alignments of the same genomic region using 4-channel images. Each color channel (here depicted using the Cyan Magenta Yellow blacK color model) encodes the following information, in order, (i) *aligned* portions for the first sample, (ii) *reference skip* portions for the first sample, (iii) *aligned* portions for the second sample, and (iv) *reference skips* portions for the second sample. This example region exhibits an *Exon Skipping/Cassette Exon* event.

Since maximum coverage and length of the region generally vary across the regions of interest, the images were resized to a width of 1000 pixels and height equal to 200, 400, 800 depending of the encoding before being fed to the network that, by definition, requires same-sized inputs.

### 2.2 Generation of training images

Training images have been generated using synthetic RNA-seq samples to ensure to obtain a comprehensive and curated set of labeled examples, necessary for supervised learning. We started from the GRCh38 v109 gene annotation keeping only protein-coding genes with multiple multi-exon transcripts annotated with the Havana/Havana TAGENE pipeline. Then, from the filtered annotation we extracted the regions of interest by pairwise comparison of the transcripts of the same gene and keeping all the regions where an AS event, as defined in [15] between the two transcripts is present. For each pair of transcripts exhibiting a local AS event, we simulated RNA-seq reads using Polyester [28] and we aligned them to the GRCh38 reference genome using STAR [29]. For each encoding defined in Section 2.1, we then generated 4 images for each pair *t*_*i*_, *t*_*j*_ of transcripts and for each region of interest involving *t*_*i*_ and *t*_*j*_ by considering (1) the alignments of the reads simulated from *t*_*i*_ as sample 1 and the alignments of reads simulated from *t*_*j*_ as sample 2, (2) the opposite, *i*.*e*., reads of *t*_*i*_ as sample 2 and reads of *t*_*j*_ as sample 1, (3) reads of *t*_*i*_ as both sample 1 and sample 2, and (4) reads of *t*_*j*_ as both sample 1 and sample 2. Cases (1)–(2) correspond to the presence of the AS event between the two samples, and the training images generated from these cases are labeled with the corresponding local AS event they exhibit, using the following nomenclature: *A* (Alternative Acceptor/Donor sites), *CSE* (Exon Skipping/Cassette Exon), and *RI* (Intron Retention). On the other hand, cases (3)–(4) correspond to the *absence* of AS events between the two samples, and the images generated from these two cases are used to train the model to classify also the *non-event* (labeled *NN*).

We have added complexity to the training images, so that those images better resemble real data. More precisely, we generated some cross-fed samples, that is we have randomly selected a portion of the reads from one sample and we have added those reads to the other sample. Such operation yields two interesting and useful results; (i) it adds noise to the images, making the network more robust to external noise and (ii) it simulates cases in which both the two different splicing events are concurrently present, albeit at different expression levels. In our experiments, we have a 20% cross-feed in both directions. An example of the images resulting from this process is shown in Figure 4.

**Figure 4:**
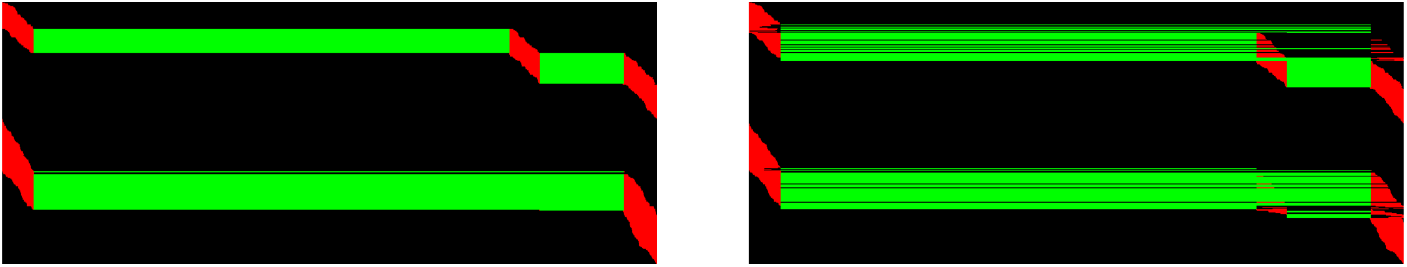
Cross-feed example: (left) each sample expresses a single transcript, (right) same region with a cross-feed of 20% in both directions. Images generated by the *Squish* encoding method.

### 2.3 Deep Learning model architecture

In order to perform region classification within our study, we employed a residual neural network architecture known as ResNet50 [30]. Since our datasets comprises 2-color-channel images, and 4-color-channel images, an adapted version of ResNet50 for each case was used. This adaptation of the ResNet50 architecture only considers the change in the input size to handle both input images, *i*.*e*.intermediate layers remain unaltered.

The last layer of the neural network is a dense layer with *softmax* activation function, to assign a single class label to each region under consideration. To train our model, we selected the *categorical crossentropy* loss function.

Each model was individually trained for 10 epochs with a batch size of 8 using *Stochastic Gradient Descend* with a learning rate of 0.001 and momentum of 0.9 [31]. The validation loss was monitored after each epoch to retain the overall best training weights.

For the classification of real samples, all trained models are ensembled. The final classification is performed using a *soft-voting* procedure, in which each network emits its confidence score for every class and the softmax of the sum of all models is then used for the final prediction.

### 2.4 Training the model

Each model has been trained on its respective image representations starting from the same set of annotated regions. Each dataset of images consisted of approx 57000 images which was then randomly split into training and test set with a ratio of 70:30.

Firstly, we performed a 5-fold cross-validation by randomly splitting the training set into training and validation and evaluating each fold against the entire test set, with the goal of testing the absence of overfitting and bias selection and to ensure the ability of classify new data. Table 1 shows the average and standard deviation of precision, recall and F1-score across the 5 folds; where `deepSpecas` clearly express extremely good classification results, with all measures having a score greater than 99%.

**Table 1:** Accuracy scores for 5-fold cross-validation show an elevated accuracy of the model on all the image representations, achieving *>* 99% on all measures.

| Method | Precision (5-fold) | Recall (5-fold) | F1-score (5-fold) |
| --- | --- | --- | --- |
| Cov | $0.9964 \pm 0.0015$ | $0.9906 \pm 0.0058$ | $0.9934 \pm 0.0036$ |
| Squish | $0.9975 \pm 0.0006$ | $0.9945 \pm 0.0022$ | $0.9960 \pm 0.0013$ |
| Comb | $0.9955 \pm 0.0016$ | $0.9852 \pm 0.0023$ | $0.9902 \pm 0.0011$ |
| Cov-4D | $0.9990 \pm 0.0004$ | $0.9989 \pm 0.0004$ | $0.9989 \pm 0.0004$ |
| Squish-4D | $0.9987 \pm 0.0009$ | $0.9968 \pm 0.0019$ | $0.9978 \pm 0.0013$ |
| Comb-4D | $0.9994 \pm 0.0006$ | $0.9994 \pm 0.0004$ | $0.9994 \pm 0.0005$ |

We then retrained the models on the entire (sub-sampled) training set for further analysis; the results of the entire test set for each model are shown in Figure 5 as confusion matrices and all the models achieve an F1-Score greater than 99%.

**Figure 5:**
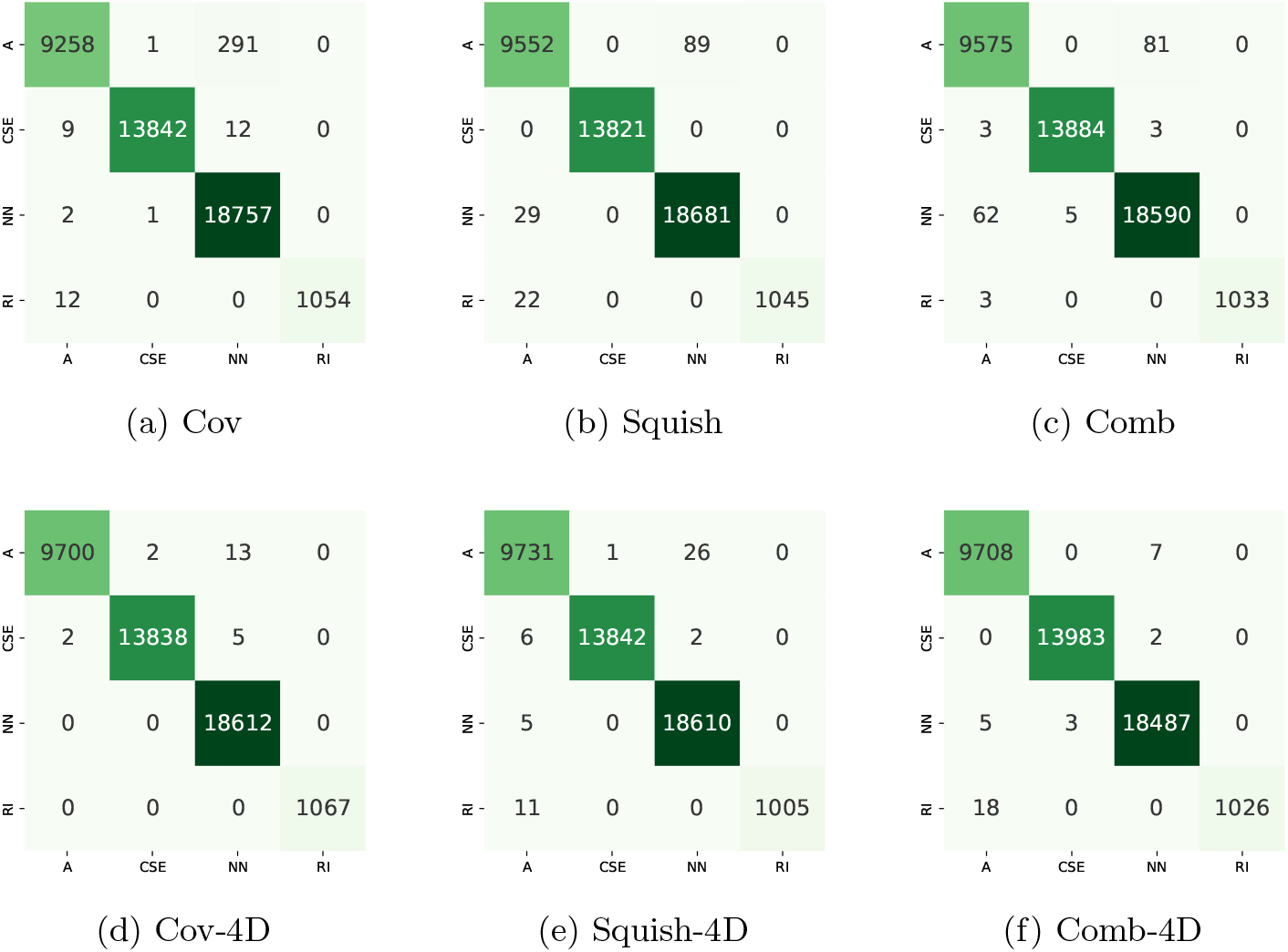
Confusion matrices for all the models obtained on the entire test set. All the models achieve an F1-score *>* 99%.

## 3 Results

### Dataset

To assess the efficacy of `deepSpecas` in classifying AS events, we performed an experimental evaluation on real RNA-Seq data. To this aim, we considered the dataset provided in [32] (SRA BioProject ID: PRJNA255099) and extensively used to evaluate tools for the analysis of AS events [15, 12]. In the study, control samples are compared with samples obtained after the double knockdown of the TRA2 splicing regulatory proteins (TRA2A and TRA2B).

The dataset consists of six Illumina Hiseq2000 samples (three replicates for two conditions) of 101bp-long paired-end reads. A total of 81 exon skipping events have been RT-PCR validated in this control-vs-knockdown dataset.

Since we want to assess also the ability of `deepSpecas` to detect other AS events, we have extended the ground-truth set of events besides the RT-PCR validated exon skipping events, by adding all AS events that are reported and considered significant (*i*.*e*., events with reported *p*-value *<* 0.05) by both rMATS [9] and SUPPA2 [15] — two state-of-the-art tools for differential analysis of AS events. Overall, the ground-truth set of AS events has 62 alternative acceptor (A3), 54 alternative donor (A5), and 41 intron retention events, in addition to the 81 RT-PCR validated exon skipping events.

We have manually inspected the ground truth set and we have found several cases where the consensus of rMATS [9] and SUPPA2 [15] is likely to be incorrect or uncertain — we detail in Figures 6 and 7 two such cases and the rationale for our decision on whether to exclude each of them from the ground truth. First consider Figure 6, it is evident that the second condition (in green) contains a high amount of reads expressing an exon skipping compared to the first condition (in red). Although there are different annotated transcripts and it is affected by spuriously aligned reads, there clearly is a different event expressed by the two conditions.

**Figure 6:**
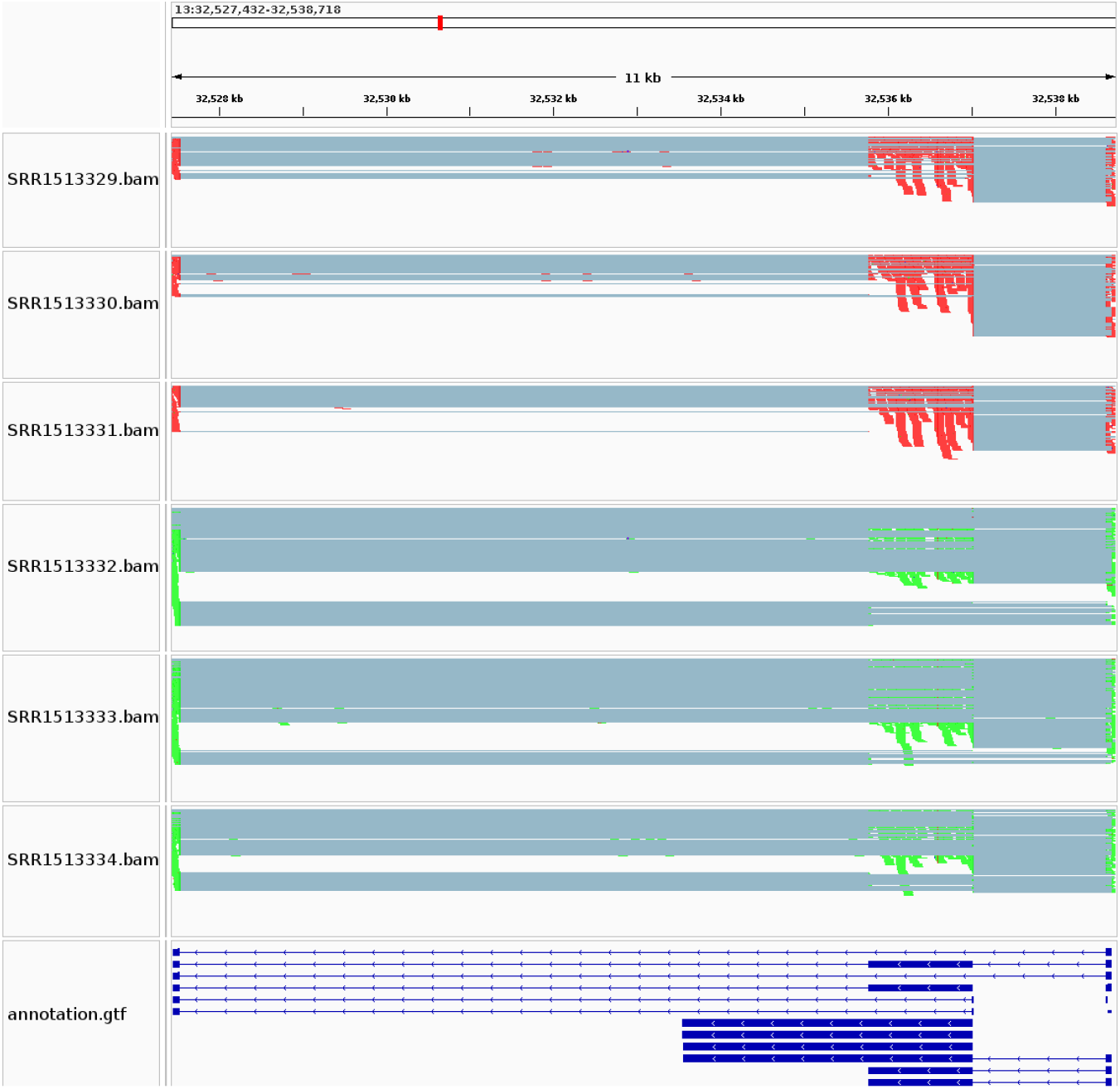
Example of an exon skipping event reported by both `SUPPA2` and `rMATS` as differentially expressed in the two samples (indicated in red and green). Even if spurious reads are present, it is evident that green condition expresses an exon skipping compared with the red one. This event was included in the manually-curated subset of events.

**Figure 7:**
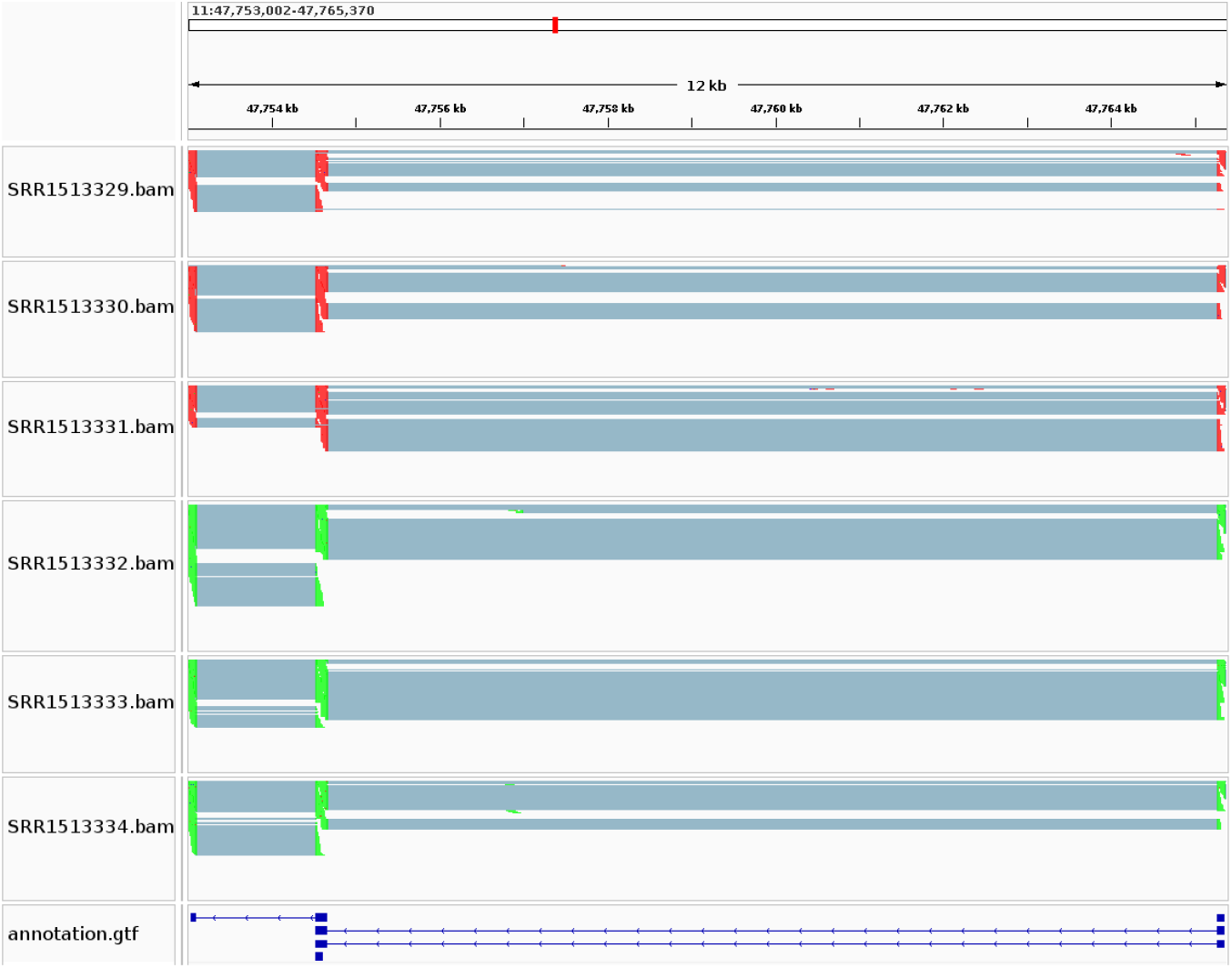
Another example of an exon skipping event reported by both `SUPPA2` and `rMATS` as differentially expressed in the two samples (indicated in red and green). However, in this case, we believe that there is not enough evidence supporting the differential expression of the splicing event. Indeed, the annotation does not include a transcript with the exon skipping event and the levels of expression in the two conditions are very similar. This event was *not* included in the manually-curated subset of events.

On the contrary the region in Figure 7 does not contain a gene annotation that support an exon skipping event, but more importantly the two conditions do not express two different events; indeed they contain roughly the same amount of reads that cover and skip the exon in the middle.

In our curation, we do not consider the transcript annotations, as they might not be complete, and we greatly prioritize the different levels of expression present in the two conditions. With this assumption, we then filtered out events that did not respect such criterion, even if RT-PCR valided (in the case of CSE) or if `rMATS` and `SUPPA2` agreed on the call.

After a manual inspection, we removed all regions in which (i) the event was supported by a very small number of reads, (ii) the region contained multiple AS events, (iii) there was no significant difference between the two samples, hence no differential expression between samples. Overall, we removed 16/81 (20%) CSE events, 34/62 (54%) A3 events, 24/54 (44%) A5 events, and 24/41 (58%) RI events.

In the following, we refer to the ground truth set before our curation as the *non-curated* dataset, while the subset obtained after our manual inspection as the *curated* dataset. The complete collection of dataset and scripts is available in the repository and the entire experimental analysis is reproducible via a provided Snakemake file. Furthermore the curated dataset is available at https://github.com/sciccolella/deepSpecas/tree/master/benchmark with the regions and the subset of reads belonging to them divided by conditions, for replication of our results and as possible benchmark dataset.

### AS analysis

We performed the AS classification on such datasets using `deepSpecas` with the *soft-voting* ensemble of all the models obtaining the results shown in Table 2 (Non-curated) where `deepSpecas` correctly classify between 70% and 80% across the different events. When we perform the same classification on the curated data, we see a considerable increase in performance of our method – CSE +2.4%, A3 +6.25%, A5 +19.2%, RI +14.5% – with respect to the non-curated as shown in Table 2 (Curated). It is interesting to notice that while we filtered all the events that we considered to not be valid we retained regions that, even if not absolutely evident, still showed some level of differential expression across the conditions, thus showing the ability of `deepSpecas` to classify noisy data. Moreover it is important to consider that `rMATS` and `SUPPA2` do call all the considered regions, but they also perform many debatable call, implying high recall at the expense of precision. Furthermore, they do rely on the gene annotation and they use it to perform the classification, while, on the other hand, `deepSpecas` does not use such information and shows considerable performance while avoiding possible biases due to the use of incomplete annotation.

**Table 2:** Results of deepSpecas classification on the real AS events datasets obtained by consensus of rMATS and SUPPA2, where we show the number of events that are correctly classified and the total number of events, for each type of AS event. The manually curated dataset shows a significant improvement over the non-curated events. Furthermore, it is interesting to notice that even in very noisy settings deepSpecas is still able to perform an acceptable classification.

|  | CSE | A3 | A5 | RI |
| --- | --- | --- | --- | --- |
| Non-curated | 67/81 (83%) | 50/62 (81%) | 42/54 (78%) | 29/41 (70%) |
| Curated | 55/65 (85%) | 24/28 (86%) | 28/30 (93%) | 14/17 (82%) |

Lastly Figure 8 shows the confusion matrices for the real datasets, in both non-curated (a) and curated (b) we can see a high accuracy of results with no particular class that overshadow the others. In both dataset no *non-event* (NN) is present as input, given their construction. Furthermore, it is also clear that a number of events that were correctly classified in the non-curated dataset are eliminated by the manual curation, thus clarifying that no bias towards `deepSpecas` was employed when filtering the original dataset.

**Figure 8:**
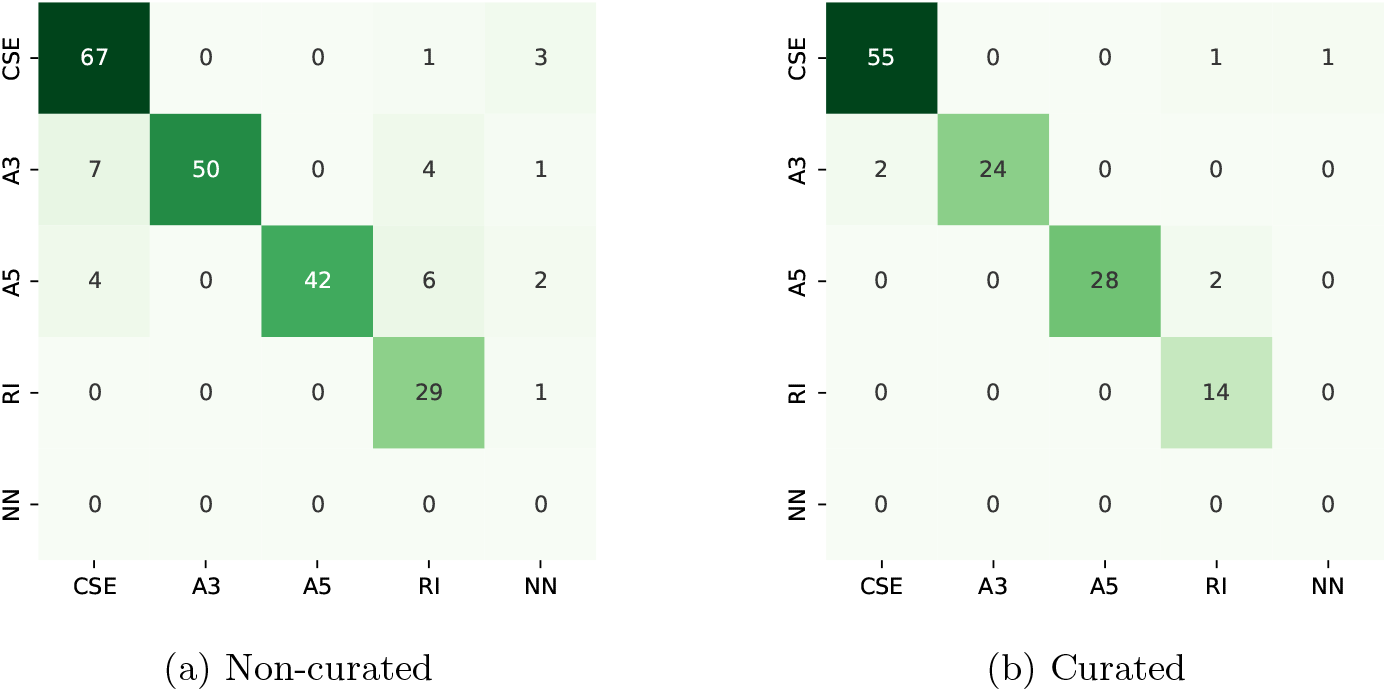
Confusion matrices of `deepSpecas` classification on the real AS events dataset obtained by consensus of `rMATS` and `SUPPA2`. No *non-event* (NN) region is present in the dataset since they are consensus of calls of the other tools (a) and furthermore a manually curated subset of them (b).

## 4 Conclusions

In this paper we devised some novel image representations of differential alternative splicing events and developed a deep neural network model trained on these dataset to accurately classify AS events. We trained a different network for each image type, achieving high accuracy levels. Furthermore, we developed a soft voting approach that leverages the different information provided by each image model, to provide a final classification by ensemble of the previous networks. We showed that `deepSpecas` achieves very good performance also on real datasets, even in the presence of highly noisy data. More importantly, `deepSpecas` does not use gene and transcript information to perform its classification step, thus avoiding any bias derived from such annotation. This lack of bias makes `deepSpecas` an interesting tool to be used when the annotation cannot be completely trusted, such as in cancerous cells, or when we have only incomplete information.

Crucially, we also created a manually curated dataset of AS events that can be used to evaluate the performance of future studies. Indeed, while there are several tools to predict AS events, the field does not have a benchmark to compare different tools. We have manually inspected a large set of predicted events, building a set of supported events which can be used for driving future improvements, of `deepSpecas` and of other tools, even though this set of events is quite small. Alongside `deepSpecas`, we also released such a benchmark of regions with the set of aligned reads, already divided by event type and ready to be used in future studies.

The current version of `deepSpecas` is designed to work on specific regions that are selected by the user, since it is not fast enough to be run on a genome-scale investigation. Future work will be devoted to extend `deepSpecas` so that it is able to predict AS events on a much large scale, such as on an entire transcriptome.

## Data Availability

All the data used in this work, the source code and pipelines to reproduce the experiments are available as open-source at the repository https://github.com/sciccolella/deepSpecas.

## Conflict of interest

The authors declare no conflict of interest.

## Funding

This research work is supported by the grant MIUR 2022YRB97K, Pangenome Informatics: from Theory to Applications (PINC).

## Notes

### Competing Interest Statement

The authors have declared no competing interest.

